# The Role of ATP in the RNA Translocation Mechanism of SARS-CoV-2 NSP13 Helicase

**DOI:** 10.1101/2021.05.21.445152

**Authors:** Ryan Weber, Martin McCullagh

## Abstract

The COVID-19 pandemic has demonstrated the need to develop potent and transferable therapeutics to treat coronavirus infections. Numerous antiviral targets are being investigated, but non-structural protein 13 (nsp13) stands out as a highly conserved and yet under studied target. Nsp13 is a superfamily 1 (SF1) helicase that translocates along and unwinds viral RNA in an ATP dependent manner. Currently, there are no available structures of nsp13 from SARS-CoV-1 or SARS-CoV-2 with either ATP or RNA bound presenting a significant hurdle to the rational design of therapeutics. To address this knowledge gap, we have built models of SARS-CoV-2 nsp13 in Apo, ATP, ssRNA and ssRNA+ATP substrate states. Using 30 *μ*s of Gaussian accelerated molecular dynamics simulation (at least 6 *μ*s per substrate state), these models were confirmed to maintain substrate binding poses that are similar to other SF1 helicases. A gaussian mixture model and linear discriminant analysis structural clustering protocol was used to identify key aspects of the ATP-dependent RNA translocation mechanism. Namely, four RNA-nsp13 structures are identified that exhibit ATP-dependent populations and support the inch-worm mechanism for translocation. These four states are characterized by different RNA-binding poses for motifs **Ia**, **IV** and **V** and suggest a powerstroke–like motion of domain 2A relative to domain 1A. This structural and mechanistic insight of nsp13 RNA translocation presents novel targets for the further development of antivirals.

## Introduction

Severe acute respiratory syndrome coronavirus 2 (SARS-CoV-2), responsible for the COVID-19 pandemic, has infected over a 150 million people and caused more than 3 million deaths worldwide as of May 2021.^1^ While antigen-based vaccines have demonstrated significant success at mitigating severe disease and spread, the need to treat infected patients as well as the evolution of potentially vaccine resistant mutants make the development of potent antivirals a pressing concern. Additionally, it has been suggested that there is the potential for new coronaviruses to become infectious to humans necessitating the development of alternative and possibly general therapeutics. To that end, the characterization of structure-function relationship of vital SARS-CoV-2 proteins is necessary to aid in the development of antivirals.

A promising target for antiviral drug development against SARS-CoV-2 is the nonstructural protein 13 (nsp13), one of 16 non-structural proteins,^2^ because it plays a critical role in viral replication and the inhibition of which in SARS-CoV-1 has been demonstrated to lead to inhibition of viral replication.^3,4^ Nsp13 is a helicase protein that is highly conserved across SARS viruses^5–7^ and is hypothesized to be a component of the RNA replication complex with the RNA polymerase, nsp12, and other nsps.^2,7–10^ Nsp13 has also been implicated in viral RNA capping activity^7,11^ and as an interferon antagonist.^12^ Viral helicases have been targets for antiviral development in SARS,^2,6,13^ flaviviruses^5,14–24^ and other positive-sense RNA viruses. Further characterization of the structure–function relationships of SARS-CoV-2 nsp13 will allow for clarification of its role in viral replication and aid in the development of antivirals.

SARS-COV-2 nsp13 is classified as a Superfamily 1 (SF1) helicase allowing for the prediction of the ATP-pocket and RNA-binding cleft within its structure. While there is no SARS-CoV-2 nsp13 crystal structure available in the literature, the very close homolog from SARS-CoV-1 (PDB: 6JYT),^25^ has a 99.8% sequence identity with SARS-CoV-2 nsp13.^7^ The SARS-CoV-1 nsp13 structure, depicted with subdomain coloring in Figure 1(b), is composed of five domains: zinc binding domain (ZBD, red), stalk domain (blue), domain 1B (pink), domain 1A (green), and domain 2A (cyan). Nsp13 has been classified as a SF1 Upf1-like helicase, a family of enzymes with similar sequence characteristics, such as distinct structural features, specificity for ATP, and unwinding polarity, that are found to interact with DNA/RNA in both eukaryotes and viruses.^25–29^ As found in all SF1 and SF2 helicases, domains 1A and 2A, also known as the RecA-like domains, comprise the conserved helicase region. The classification of nsp13 as a SF1 helicase allows for the prediction of the ATP binding site in-between domains 1A and 2A as well as the RNA-binding cleft separating the RecA-like domains from domain 1B. ATP is depicted in its predicted binding pocket in Figure 1(c). Verification of these structural inferences is necessary for developing well-founded structure–function hypotheses as well as rationally designing therapeutics.

**Figure 1.**
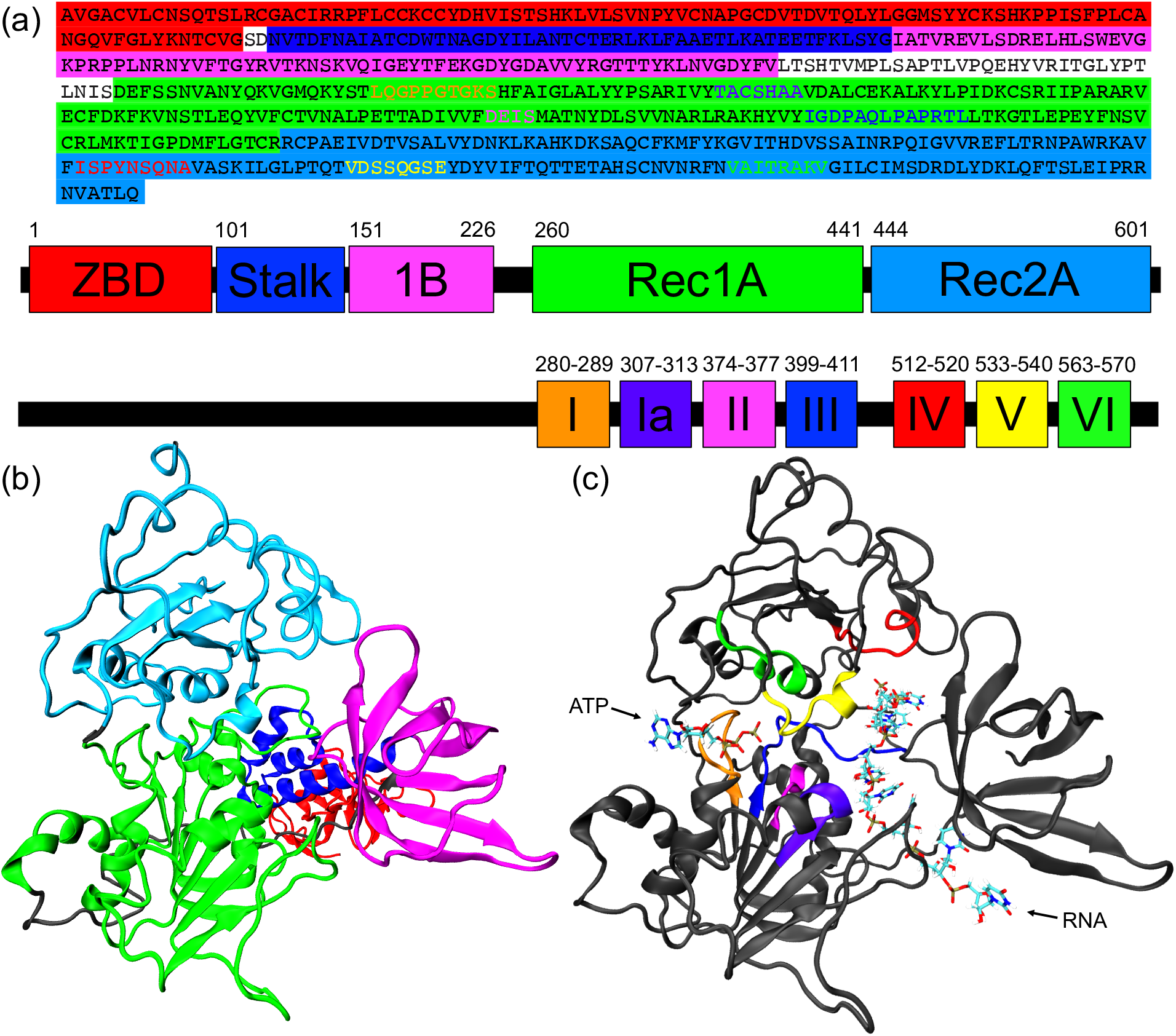
SARS-CoV-2 Nsp13 helicase structure. (a) Sequence, domain structure, and motif structure of nsp13. Nsp13 model based on the I570V mutation of SARS-CoV-1 nsp13 (6JYT), colored by (b) domain and (c) motif.^30^ The Zinc Binding Domain and Stalk Domains were removed in the motif structure for clarity.

Aspects of the structure–function relationship for nsp13 can be inferred through comparison to sequence-similar enzymes. Upf1-like helicases utilize an NTPase cycle to provide the free energy to unwind dsRNA and translocate along the nucleic acid substrate in a 5’ to 3’ direction.^25,27^ These enzymes also exhibit RNA-dependent NTPase activity. ^31^ A set of highly conserved motifs including NTP binding and hydrolysis motifs (**I**, **II**), RNA binding and unwinding motifs (**Ia**, **Ib**, **IV**), and motifs connecting the two binding regions (**III**, **V** and **VI**) have been found to be important for the function of SF1 helicases.^26–28,32^ The motifs found in SARS-CoV-2 nsp13 are indicated in Figure 1(c). How all of these motifs work in concert to allow for NTP-dependent RNA translocation remains unknown for SARS-CoV-2 nsp13, yet this information is critical for rational design of inhibitors.

SF1 helicases are thought to translocate by either an inchworm stepping or Brownian ratchet mechanism. ^33^ The inchworm mechanism has two sites that alternate between strongly and weakly bound states such that one site is always strongly bound to RNA. The weakly bound site performs a power stroke before strongly binding RNA one basepair forward. This behavior is dependent on the ATP substrate state and can lead to unidirectional translocation along an oligonucleotide substrate. The Brownian ratchet is a simpler two state model in which the RNA is either strongly or weakly bound to the protein. RNA is translocated through a power stroke when the protein enters the short lived, weakly bound state. These mechanisms are distinguished by their ATP-dependent RNA-binding activity and, as such, are testable based on ATP-dependent RNA-bound structures. The mechanism of RNA-translocation by SARS-CoV-1 nsp13 has been examined at an ensemble kinetics level^2^ and singlemolecule kinetics level^34^ to identify the rate of ATP hydrolysis and translocation. H/D mass exchange data suggest the presence of at least two RNA-bound states depending on the presence of an ATP substrate.^25^ These data are insufficient, however, to distinguish between the two proposed mechanisms and do not provide enough structural information to inform antiviral development.

The work presented here examines the structure–function relationships utilized by SARS-CoV-2 nsp13 during ATP-dependent RNA translocation. Specifically, we focus on the effect of the presence of ATP and RNA on the structural ensemble of the protein and elucidate the role of ATP in the translocation mechanism. We perform extensive all-atom Gaussian accelerated molecular dynamics (GaMD)^35^ in combination with Gaussian Mixture Model (GMM) structural clustering and characterization by Linear Discriminant Analysis (LDA). We identified four states in the RNA-binding cleft which is indicative of nsp13 translocating using an inchworm stepping mechanism. The role of ATP binding in the translocation mechanism is also elucidated. Furthermore, we analyze the ATP-pocket of the four states identifying key motifs that allosterically connect the ATP-pocket to the RNA-binding cleft.

## Computational Methods

### System Setup

Simulations were performed for four ligand-bound states of the nsp13 helicase: Apo, ssRNA, ATP, and ssRNA+ATP. The initial structure for the Apo state is based on the I570V mutation of SARS-CoV-1 (PDB: 6JYT).^25^ Due to the lack of ligand-bound crystal structures of SARS nsp13s, RNA and ATP substrates were extracted from Upf1-like helicases aligned to the mutated SARS-CoV-1 nsp13 crystal structure. For the ssRNA and ssRNA+ATP states, polyuracil ssRNA was extracted from the RNA-bound Upf1 helicase crystal structure (PDB: 2XZL)^36^ after the two crystal structures were aligned using a sequence–based maximum likelihood protocol as implement in THESEUS.^37^ For the ssRNA+ATP bound state, the ATP-analog AMP-PNP and the coordinated Mg^2+^ ion were extracted from the AMP-PNP-bound Upf1 helicase crystal structure (PDB: 2GJK)^38^ after the two crystal structures were aligned using the P-loop, motif **II**, motif **VI**, and motif **V** as alignment landmarks. The amide group between the *β-* and *γ*-phosphates in AMP-PNP were replaced by an oxygen atom to form ATP. Furthermore, residues Gly282-Gly287 from the P-loop region in domain 2A were replaced by residues Gly430-Gly435 from the P-loop region in the AMP-PNP-bound Upf1 helicase. The ATP state was created by taking the ssRNA+ATP state and removing the ssRNA.

The protein is modeled using ff14SB parameters,^40^ RNA is modeled using ff99bsc0_*χOL*3_ parameters,^41,42^ and the parameterization files for ATP^43^ are obtained from the AMBER parameter database. Additionally, nsp13 has a zinc binding domain with three nonstandard zinc binding pockets. The three zincs are parameterized in Cys-Cys-Cys-Cys, Cys-Cys-Cys-HID, and Cys-Cys-HID-HIE environments, respectively, using the MCPB tool in AMBER. ^44^ Crystollagraphic waters are maintained for each state and TIP3P water was added to each system with at least a 12 A buffer yielding a cubic box of linear dimension of at least 125 Å and a total of at least ~ 175K atoms. Na^+^ and Cl^-^ ions were added to neutralize charge and provide a 0.1 M salt concentration.

### Simulation Details

All-atom, explicit solvent GaMD simulations for the Apo, ssRNA, ATP, and ssRNA+ATP states of nsp13 are performed using the GPU-enabled AMBER18 software.^39^ Hydrogen atoms are constrained using the SHAKE algorithm.^45^ Direct nonbonding interactions are cut off at 12 Å, and long-range electrostatic interactions are modeled using the particle mesh Ewald treatment (PME). ^46^ An integration time step of 2 fs is used. GaMD simulations are performed in the NPT ensemble with a Monte Carlo barostat set to 1 atm and a Langevin thermostat set to 300 K.

The simulation protocol used for all systems are the same. Systems are minimized in ten stages. In all minimization stages 2000 steepest descent minimization steps are performed with varying harmonic restraints. In the first stage there is a 500 kcal mol^−1^ Å^−2^ restraint on all protein and ligand atoms. In the next four stages the restraints on the protein sidechains are reduced to 10.0, 1.0, 0.1, and 0.0 kcal mol^−1^ Å^−2^, respectively. Finally, in the last five stages there are diminishing restraints on ATP+RNA the protein backbone and all ligands of 50.0, 5.0, 1.0, 0.1 and 0.0 kcal mol^−1^ Å^−2^. The system is heated to a temperature of 300K over 1 ns with a harmonic restraint of 40 kcal mol^−1^ Å^−2^ on all protein and ligand atoms. Pressure equilibration is performed in six stages. First 1 ns of NVT simulation is performed maintaining the harmonic restraint from the heating. In the next five pressure equilibration stages the restraint was reduced to 20.0, 10.0, 5.0, 1.0, and 0.1 kcal mol^−1^ Å^−2^ for 200 ps each. A conventional MD (cMD) NPT simulation is then performed for 10 ns.

Following the cMD equilibration process each substrate state is simulated using GaMD. For GaMD simulations, the threshold energy for applying boost potential is set to E = Vmax and the default 6 kcal mol^−1^ is used for both *σ*_0*P*_ and *σ*_0*D*_. The maximum, minimum, average, and standard deviation values of the system potential are obtained from an initial 10 ns cMD simulation with no boost potential. Then GaMD simulations are performed with boost potential applied to both the dihedral and total potential energy terms. Each GaMD simulation is proceeded with a 40 ns equilibration run after adding the boost potential, followed by 2 *μ*s of production runs. The GaMD simulations of all substrate states are performed in triplicate except for the ssRNA bound state in which six replicates were performed yielding a total of 30 *μ*s of GaMD simulation to be anlayzed.

### Model Verification

Due to the lack of ATP-bound and ssRNA-bound crystal structures, the ATP, ssRNA, and ssRNA+ATP starting structures were created from combining ATP-bound and ssRNA-bound Upf1 helicase crystal structures with a I570V mutated SARS-CoV-1 nsp13 apo helicase crystal structure. The contacts between the SARS-CoV-2 nsp13 protein and the bound ATP and ssRNA ligands for the ATP, ssRNA, and ssRNA+ATP systems are compared to similar contacts in other SF1 helicase proteins, including the Upf1 and IGHMBP2 helicases, to show that these are suitable initial structures that are stable during simulation. For the RNA-bound systems, contacts between motifs **Ia**, **IV**, and **V** with ssRNA phosphates are determined for each frame as these motifs are highly conserved across SF1 helicase proteins. Similarly, the contacts between motifs **I**, **II**, **III**, **V** and **VI** with ATP and Mg^2+^ are calculated in the ATP-bound systems. A residue and ligand were considered to be in contact if any atom of the residue is within 5 A of any atom in the ligand. The residue identities and the percentage of frames that each residue is in contact with the ligand are shown in the supporting information for both the ssRNA and ATP contacts in Table S1 and Table S2, respectively. These tables show that a majority of the contacts formed in the Upf1 and IGHMBP2 crystal structures are maintained for 60-100% of the frames in the simulation, although, some of the contacts are made or broken depending on whether both or only a single ligand are bound to the protein.

### Gaussian Mixture Model Clustering and Linear Discriminant Analysis

Variational Bayesian Gaussian mixture model is a probabilistic model that effectively fits a given set of data to a specified number of Gaussian distributions with unknown parameters assigning each data point to a cluster. GMM is used to identify structural states in the nsp13 protein by clustering a set of distance calculated from the nsp13 simulations using a GMM tolerance of 10^−6^. The number of clusters that the data is fit to is determined by calculating the silhouette, Calinksi-Harabasz (CH), and Davies-Bouldin (DB) scores for a range of cluster sizes from two clusters to ten clusters. Then based on maximums in the silhouette and CH scores and minimums in the DB score the number of clusters is chosen.

Linear Discriminant Analysis is a classification and dimensionality reduction tool. LDA is a supervised algorithm (data must already be clustered) that finds the linear combination of features, LD eigenvectors, that maximize cluster separation. The eigenvector with the largest eigenvalue provides the largest cluster separation. LDA is used to identify the distances that best differentiate the four states identified in the RNA-binding cleft.

The GMM clustering and characterization using LDA is performed iteratively Each iteration uses the projection of the distance data onto the previous iterations LD eigenvectors as the input data for the GMM clustering. The distance data is still used as the input data for LDA. The cycle is iterated until the distance between the current iterations projected distances and the previous iterations projected distances is below a threshold value of 10^−3^. Both GMM and LDA are performed using the machine-learning Python library scikit-learn. ^47^

## Results and Discussion

The translocation mechanism of the SARS-CoV-2 nsp13 helicase is likely an important component of the viral lifecycle and yet structural details regarding this mechanism are lacking.^10^ Here, we present a set of simulations for the Apo, ATP, ssRNA, and ssRNA+ATP bound states to provide insight into the translocation mechanism of nsp13 along ssRNA. First, we discuss large scale changes in domains 1A, 2A, and 1B between all four systems. We identify changes in the RNA-binding cleft due to the binding of ATP by analyzing differences in the ssRNA and ssRNA+ATP systems which provide insight into the translocation mechanism of SARS-CoV-2 along ssRNA. Finally, we discuss allostery between the ATP-pocket and the RNA-binding cleft focusing on how the presence of ATP changes the ATP-pocket and how those changes affect the RNA-binding cleft.

### Large-Scale Changes to Protein Structure

The interdomain distances between domains 1A, 2A, and 1B, depicted in Figure 2, are calculated to investigate large-scale changes within the nsp13 protein structure due to the presence of ssRNA and ATP. The center-of-mass of the *β*-sheets for each domain are used to calculate the interdomain distances due to their rigidity within each domain. The ATP-pocket sits at the boundary between domains 1A and 2A. The average 1A–2A domain distance remains constant around 31 Å, independent of the presence of ATP and ssRNA as demonstrated by the average separation distances shown in Table 1. The standard deviation of all distances are calculated from the set of average reweighted distance values of each replicate of the system.

**Figure 2.**
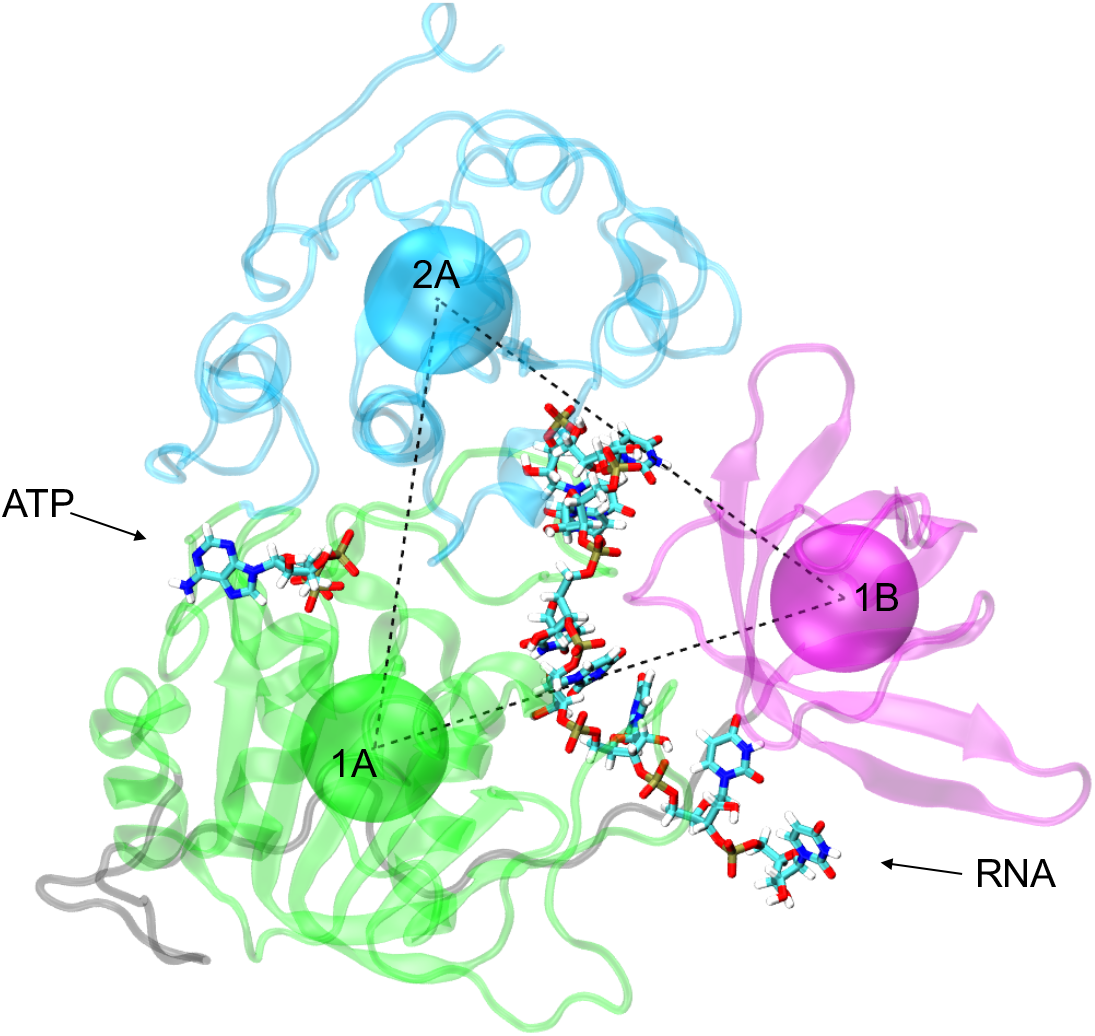
Structural depiction of the distances between the center-of-mass of domains 1B (magenta), 1A (green), and 2A (cyan). The stalk and ZBD domains are not depicted for clarity.

**Table 1.** Average center-of-mass separation distance between domains 1A, 2A, and 1B of the nsp13 helicase Apo, ATO, ssRNA, and ssRNA+ATP ligand bound states. Error in the last digit is provided in parentheses.

| Domains | Separation Distance (Å) |  |  |  |
| --- | --- | --- | --- | --- |
|  | APO | ATP | ssRNA | ssRNA+ATP |
| 1A-2A | 31.1(6) | 31.6(2) | 31(1) | 31.3(4) |
| 1A-1B | 34.6(6) | 34(1) | 36.6(8) | 38.4(4) |
| 2A-1B | 30(1) | 31.1(4) | 33(2) | 36.4(1) |

Although there are no large scale changes to the ATP-pocket, the presence of ATP leads to large scale changes to the RNA-binding cleft. Domain 1B runs along the edge of both domains 1A and 2A forming the RNA-binding cleft. The interplay of these two boundaries are important in the translocation mechanism of nsp13. The 1A–1B and 2A–1B domain distances provide insight into how these boundaries change with ssRNA and ATP binding. The 1A–1B domain distance increases when ssRNA is bound to nsp13. Furthermore, the binding of ATP into the ssRNA-bound state leads to additional widening of the 1A–1B distance from 36.6 A in the ssRNA system to 38.4 A in the ssRNA+ATP state. Similar behavior is observed between domains 2A and 1B where the presence of ATP leads to an increase in the 2A–1B distance by 3.4 A relative to the RNA-bound state. There is significantly more fluctuation in the 2A–1B distances relative to 1A-1B distances, which can be attributed to large fluctuations in the tertiary structure of the 2A domain. The widening of the RNA-binding cleft for both the 1A–1B and 2A–1B boundaries suggests a possible reduction in the binding strength of RNA within the RNA-binding cleft due to the presence of ATP. The linear interaction energy between each phosphate and the surrounding protein residues and the RMSF of each phosphate were calculated to determine the binding strength between nsp13 and ssRNA and can be found in Table S3 and Table S4 of the supporting information. In both analyses the fluctuations in these values were too large to differentiate between the RNA and RNA+ATP bound states. Therefore, it is necessary to form a more detailed description of the RNA-binding cleft for each of these systems.

### Structural Changes of the RNA-Binding Cleft due to the Presence of ATP

The presence of ATP leads to large scale changes in the RNA-binding cleft as shown by the increase in the 1A–1B and 2A–1B interdomain distances. Motifs **Ia** and **IV** are both highly conserved regions in SF1 helicases and it has been suggested that these motifs are involved in RNA binding.^30,48^ The structure of the RNA-binding cleft from each frame of these simulations were clustered based on the distances between motif **Ia**, motif **IV**, and the closest RNA phosphates (**P**) using a GMM-LDA approach. Based on these three distances, the GMM analysis separated the ssRNA and ssRNA+ATP structures into four states: **S1**, **S2**, **S3**, and **S4**. The specific residues used to calculate the distance between motifs is shown in Table S5 of the supporting information. A representative structure of the RNA-binding cleft for each state is shown in Figure 3.

**Figure 3.**
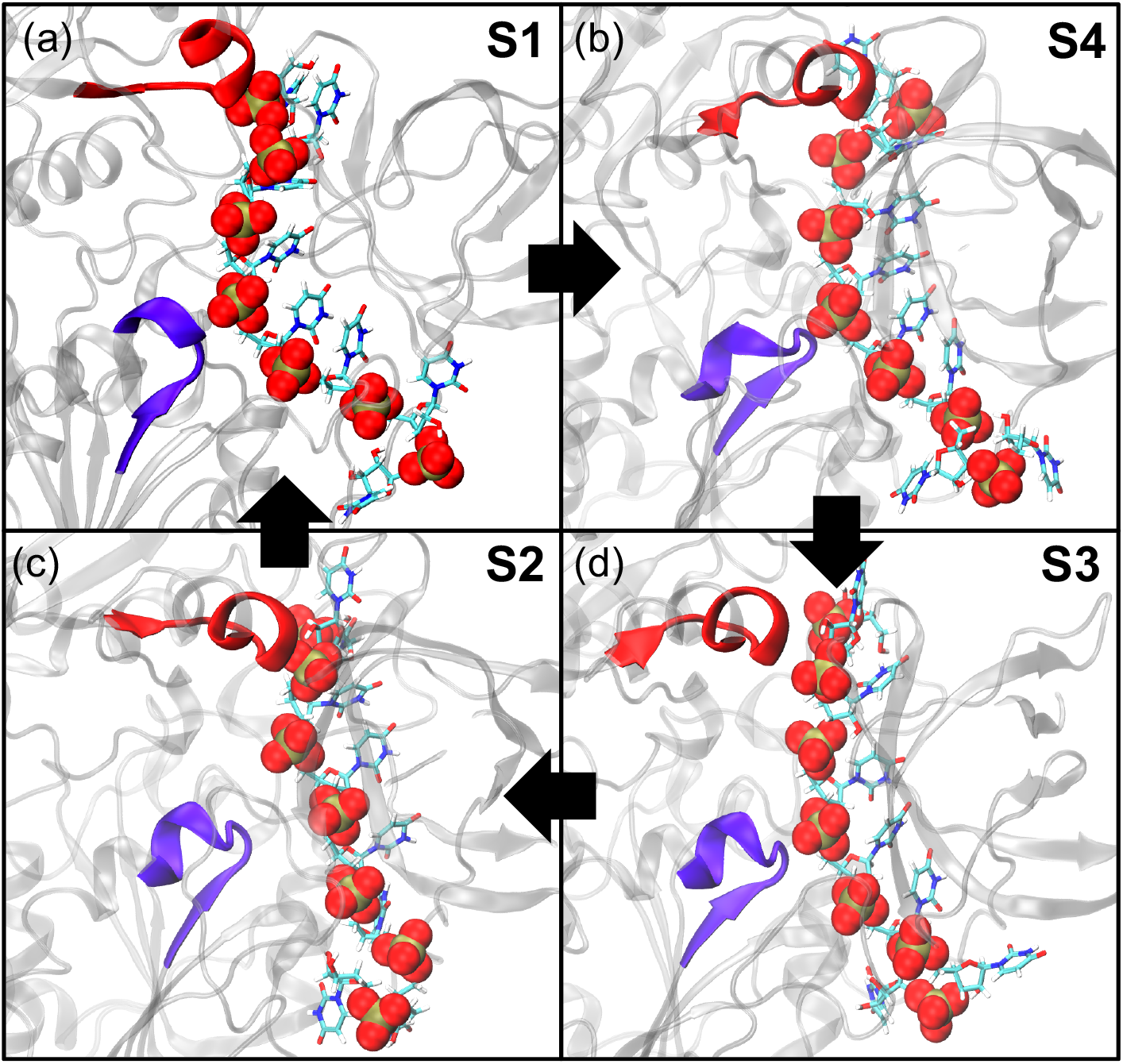
Representative structures of the states (a) **S1**, (b) **S4**, (c) **S2**, and (d) **S3** determined by the GMM-LDA clustering analysis performed on distances between motif **Ia**, motif **IV**, and the closest ssRNA phosphates. The arrows represent a 4-steps inchworm stepping translocation mechanism.

Analysis of the structure of the RNA-binding cleft for **S1**, **S2**, **S3**, and **S4** reveals that motif **Ia** and motif **IV** independently bind and release ssRNA at different phosphate positions. To determine how the structure of RNA-binding cleft varies for each of the states, the three distances were projected onto the LD eigenvectors calculated by the LDA. The coefficients of LD1 and LD2 are shown in the supporting information in Table S6. Figure 4(a,b) show the projection of the three distances on to the LD1 and LD2 eigenvectors for the RNA and RNA+ATP systems, respectively. LD1 separates **S2** from the other three states and is dominated by the distance between motif **Ia** and the RNA phosphates. Table 2 shows the average of each of the distances used in the clustering analysis for all clusters. **S1**, **S3**, and **S4** have an average **Ia** - **P** distance around 5.5 A, while in **S2** this distance increases to 9.4 A. This can be seen in Figure 3(c) where the ssRNA has separated from motif **Ia**, causing ssRNA to become more linear in the RNA-binding cleft. LD2 distinguishes between **S1**, **S3**, and **S4** and is dominated by both the **Ia** - **IV** and **IV** - **P** distances. The distance between motif **Ia** and **IV** is smallest for **S3** at 14.1 A and largest for **S4** and **S1** at 18.2 and 20.2 A, respectively. The change in the **Ia** – **IV** distance between **S1** and **S3** can be explained by a change in the number of phosphates between the phosphates bound by motifs **Ia** and **IV**. In the representative structure for **S3** (Figure 3(d)), motif **Ia** is bound to P4 and motif **IV** is bound to P2 leaving only a single phosphate gap (P3) between them. On the other hand, in the representative structure for **S1** (Figure 3(a)), motif **Ia** is bound to P4 and motif **IV** is bound to P_1_, leaving a 2 phosphate gap (P_2_ and P_3_) between them. **S1** and **S4** are not well separated by the **Ia** – **IV** distance, but are separated by the distance between motif **IV** and the RNA phosphates. In **S1**, the average **IV** – **P** distance is 5.0 Å, while in **S4** the average distance increases to 8.1 A as the ssRNA bends away from motif **IV**.

**Figure 4.**
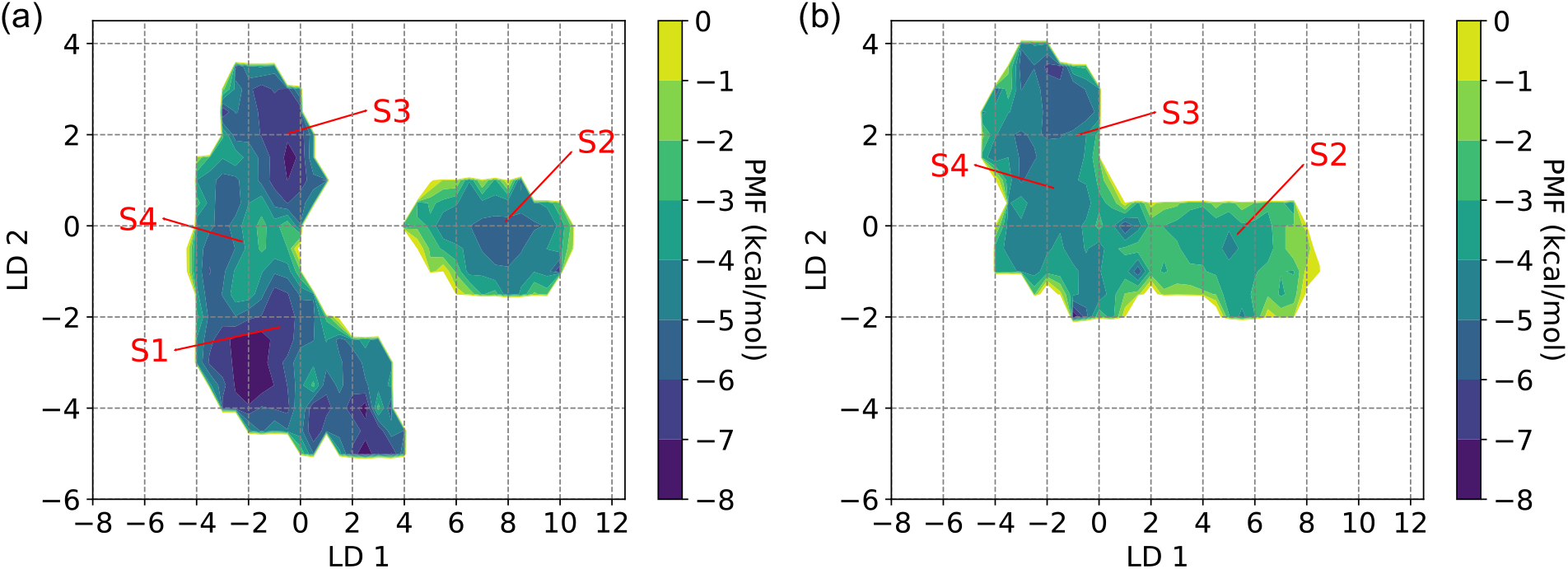
The free energy of the projection of the distances between motif **Ia**, motif **IV**, and the closest ssRNA phosphates on to LD1 and LD2 eigenvectors from the GMM-LDA clustering analysis for the (a) ssRNA and (b) ssRNA+ATP systems.

**Table 2.** Average separation distances and standard deviations between motif IV, motif Ia, and the closest ssRNA phosphates (P) for states S1, S2, S3, and S4.

| Residues | Average Distance (Å) |  |  |  |
| --- | --- | --- | --- | --- |
|  | S1 | S2 | S3 | S4 |
| **IV – P** | 5.0(8) | 4.7(4) | 6(1) | 8(1) |
| **IV – Ia** | 20(1) | 16.3(7) | 14.2(9) | 18(2) |
| **Ia – P** | 5.5(5) | 9.4(9) | 5.4(2) | 5.4(4) |

The four states identified in the GMM-LDA analysis suggest a 4-step inchworm stepping translocation mechanism of nsp13 along ssRNA. In this inchworm mechanism, motif **Ia** and motif **IV** act as binding sites that independently bind and release the phosphates of ssRNA. There is always one of the binding sites that is strongly bound to ssRNA, unlike in a Brownian rachet mechanism in which the protein as a whole, strongly or weakly binds ssRNA. The mechanism follows the cycle shown in Figure 3. Let’s assume the cycle starts in **S1** where motif **IV** is bound to P_*n*_ and motif **Ia** is bound to P_*n*+3_ leaving a two phosphate gap between them. In the first step of the mechanism motif **IV** and P_*n*_ unbind as the protein transitions to **S4**. In the second step motif **IV** does a power stroke moving down one base of ssRNA and binds P_*n*+1_ resulting in the protein being in **S3**. This leaves a single phosphate gap between the phosphates bound by motifs **Ia** and **IV**. In the third step the protein transitions to **S2** as motif **Ia** and P_*n*+3_ unbind. Finally, in the fourth step, **Ia** performs a power stroke moving down one base of ssRNA and binds to P_*n*+4_ and the protein ends up back in **S1**. Each 4-step cycle performed by nsp13 results in a one base pair translocation along ssRNA.

Changes in the sampling of **S1**, **S2**, **S3**, and **S4** when ATP is present in the ATP-pocket suggests that the binding of ATP leads to the release of ssRNA by motif **IV**. In Table 3, the probability of being in each of the four state for the ssRNA and ssRNA+ATP systems is calculated to provide insight into which of the four states the protein is in when RNA is bound and how the sampling of the four states change when ATP binds into the RNA-bound protein. In the ssRNA system, both **S1** and **S4** are each sampled 33% of the time. With ATP bound, **S1** is no longer sampled and the sampling of **S4** and **S3** both increase to 51% and 33%, respectively. These probabilities suggest that the cycle begins with a two phosphate gap between the phosphates bound by motifs **Ia** and **IV**. The binding of ATP then increases the probability of states where motif **IV** unbinds (**S4**) and performs a power stroke leading to a single phosphate gap such between motifs **Ia** and **IV** (**S3**). These probabilities provide evidence that the binding of ATP into the RNA-bound state state of nsp13 causes the first step in the inchworm stepping translocation mechanism. The remaining translocation steps most likely occur in the later steps of the hydrolysis cycle such as ATP hydrolysis and ADP and Pi release. This is supported by several studies that suggest the hydrolysis of ATP to ADP+Pi and the release of these products are responsible for the unwinding of dsRNA and leads to the power stroke resulting in the translocation of RNA helicase proteins. ^29,33,49,50^ It is necessary to perform simulations of ssRNA+ADP+Pi and ssRNA+ADP ligand bound systems to further elucidate the translocation mechanism of nsp13 along the ATP-hydrolysis cycle.

**Table 3.** The probability of being in each of the states identified in the GMM-LDA clustering analysis for both the ssRNA and ssRNA+ATP systems.

| Residues | Probability |  |  |  |
| --- | --- | --- | --- | --- |
|  | **S1** | **S2** | **S3** | **S4** |
| RNA | 33.01% | 16.45% | 15.14% | 35.39% |
| RNA+ATP | 0.02% | 15.99% | 33.22% | 50.78% |

### Structural Changes of the ATP-pocket due to the Presence of ATP

The binding of ATP to nsp13 leads to a change in the structure of the RNA-binding cleft as shown by the change in sampling of the four states identified in the RNA-pocket and the increase in the 1A–1B and 2A–1B interdomain distances when ATP is bound into the ssRNA state. To identify how the presence of ATP changes the ATP-pocket and how these changes allosterically alter the RNA-binding cleft, LDA was performed on states **S1**, **S2**, **S3**, and **S4** using distances between ATP-binding motifs (**I**,**II**,**VI**), RNA-binding motifs (**IV**), and motifs that bind both ATP and RNA (**Ia**,**V**,**III**). Initially, all residues in all conserved motifs were included in the LDA analysis. Iteratively, LDA was performed on these distances and the distances that received low coefficients or contained similar information to other distances in LD1 and LD2 were removed until the eight distances shown in Table 4 were chosen.

**Table 4.**
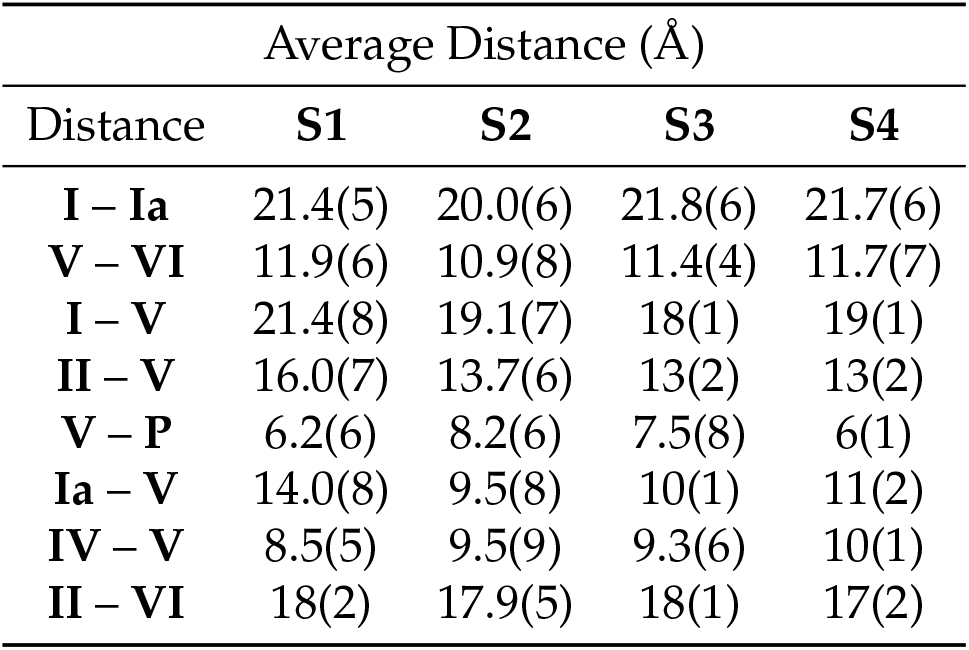
Average separation distances and standard deviations between motifs I, motif Ia, motif II, motif IV, motif V, motif VI, and the closest ssRNA phosphates for states S1, S2, S3, and S4.

| Distance | Average Distance (Å) |  |  |  |
| --- | --- | --- | --- | --- |
|  | S1 | S2 | S3 | S4 |
| I – Ia | 21.4(5) | 20.0(6) | 21.8(6) | 21.7(6) |
| V – VI | 11.9(6) | 10.9(8) | 11.4(4) | 11.7(7) |
| I – V | 21.4(8) | 19.1(7) | 18(1) | 19(1) |
| II – V | 16.0(7) | 13.7(6) | 13(2) | 13(2) |
| V – P | 6.2(6) | 8.2(6) | 7.5(8) | 6(1) |
| Ia – V | 14.0(8) | 9.5(8) | 10(1) | 11(2) |
| IV – V | 8.5(5) | 9.5(9) | 9.3(6) | 10(1) |
| II – VI | 18(2) | 17.9(5) | 18(1) | 17(2) |

Motif **V** is identified as a key motif for allosteric communication between the ATP-pocket and the RNA-binding cleft due to changes in its positioning between them for the four states identified in the GMM-LDA analysis of the RNA-binding cleft. To determine how the structure of the ATP-pocket varies in the four states, the eight distances are projected onto the LD1 and LD2 eigenvectors for the ssRNA and ssRNA+ATP systems as shown in Figure 5(a,b), respectively. The coefficients of LD1 and LD2 are shown in the supporting information in Table S7. LD1 separates **S2** from **S1**, **S3**, and **S4**. The average distances shown in Table 4 are ordered based on the LD1 coefficients. **S2** has a smaller **I** - **Ia** distance relative to the other states as domain **Ia** unbinds ssRNA and sits close to the ATP-pocket. Similarly, motif **V** is closer to both the ATP-pocket and motif **VI** for **S2**, leading to the largest **V** - **P** distance and smallest **V** - **VI** distance of all four states. Furthermore, motif **Ia** and **V** sit much closer together relative to the other states. Overall, **S2** has a much more compact ATP-pocket which leads to motifs **V** and **Ia** not binding or weakly binding ssRNA. LD2 differentiates between **S1**, **S3**, and **S4** and has the largest contribution from the and **I** - **V** distance. The **I** - **V** distance is smallest for **S3** at 18 A as motif **V** sits closer to the ATP-pocket as shown in Figure 6(c). The **I** - **V** distance increases for **S4** to 19 A and increases again for **S1** to 21.4 A as motif **V** moves closer to the RNA-binding cleft and binds to the ssRNA phosphates. This is further shown by the decrease in the **V** - **P** distance for **S1** and **S4**. Not only does motif **V** sit closer to the RNA-binding cleft for these two states, but it rotates up away from motif **Ia** becoming more parallel with the ssRNA backbone as it binds one or two of the phosphates as shown in Figure 6(a,b). This is evidenced by the increasing **Ia** – **V** and **II** – **V** distances.

**Figure 5.**
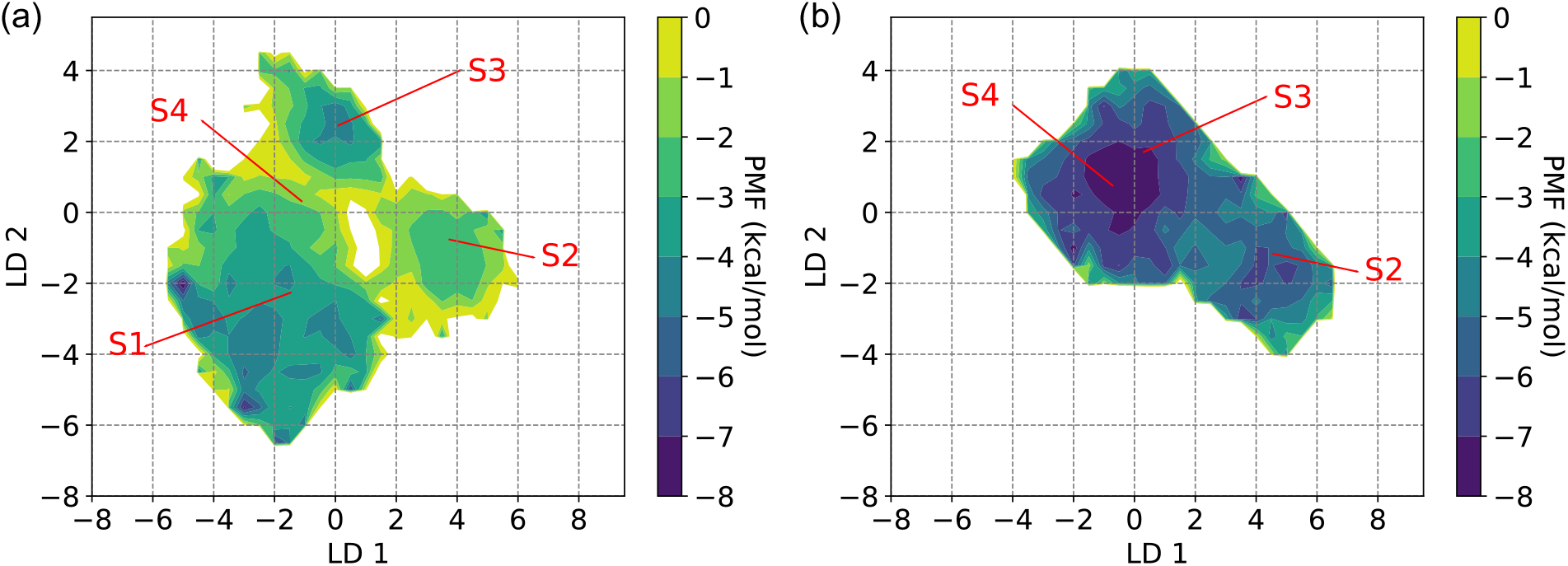
The projection of select distances between motif **I**, motif **Ia**, motif **II**, motif **IV**, motif **V**, motif **VI**, and the closest ssRNA phosphates on to LD1 and LD2 eigenvectors from the GMM-LDA clustering analysis for the (a) ssRNA and (b) ssRNA+ATP systems.

**Figure 6.**
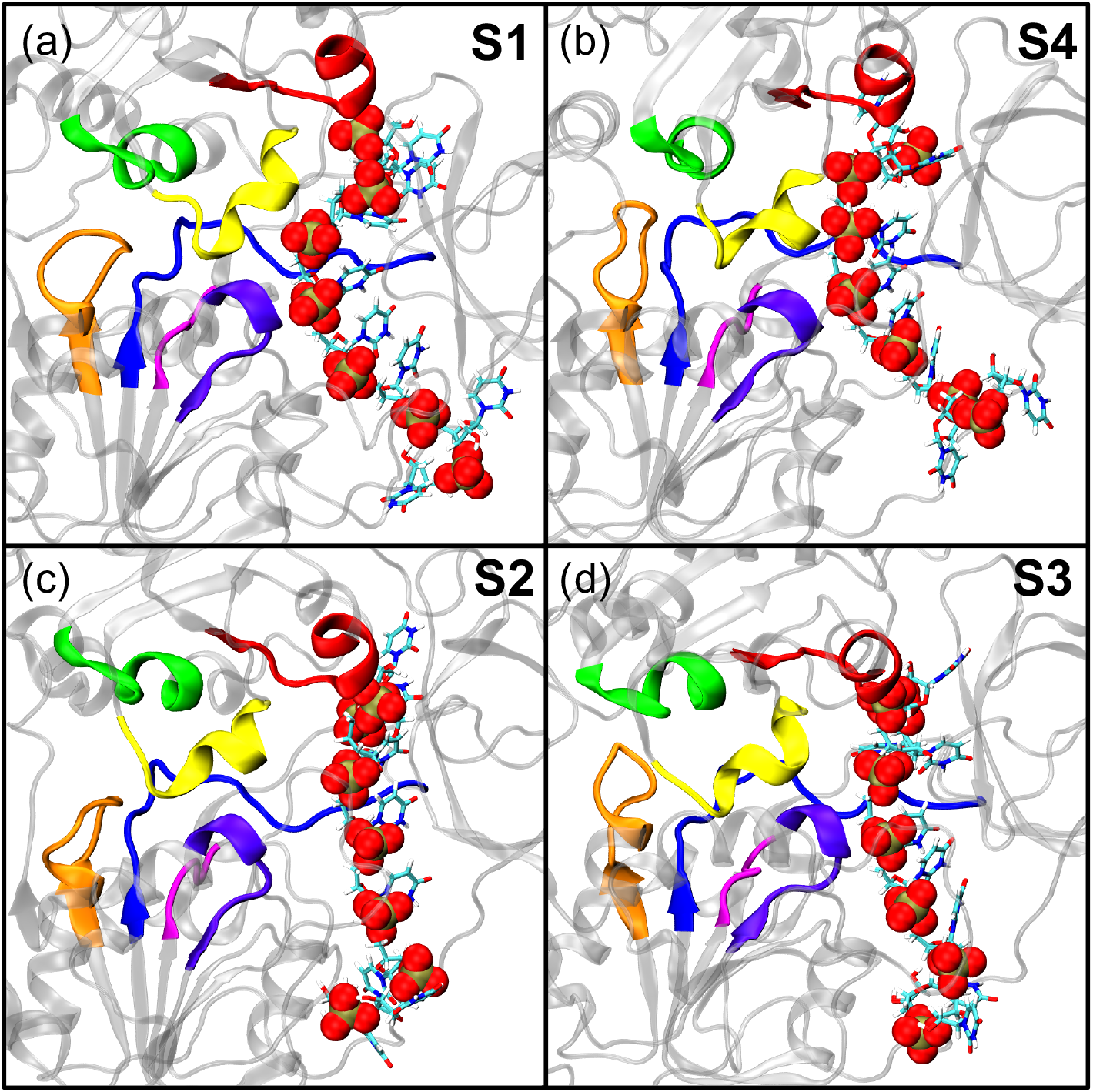
Representative structures of the ATP-pocket for states (a) **S1**, (b) **S4**, (c) **S2**, and (d) **S3** determined by the GMM-LDA clustering analysis performed on the RNA-binding cleft.

The analysis of the ATP-pocket allows us to form a more complete picture as to how the presence of ATP leads to a change in sampling of **S1**, **S2**, **S3**, and **S4**. As motif **V** moves farther from the ATP-pocket it binds to ssRNA phosphates closer to motif **IV** stabilizing the two phosphate gap between motifs **Ia** and **IV** and, therefore, stabilizing **S1**. As nsp13 binds ATP there is an increase in the sampling of **S4** and **S3** where motif **V** sits closer to the ATP-pocket due to the formation of contacts between ATP and motif **V**. Table 5 shows the average distance between all residues in motif **V** and ATP or Mg^2+^ for the ssRNA+ATP system. In **S2**, **S3**, and **S4** Val 533, Asp 534, Ser 535, and Ser 536 form contacts with ATP and Mg^2+^ with an average separation distance of 3-6 Å. These contacts make it energetically unfavorable for motif **V** to move close enough to ssRNA to stabilize the two phosphate gap between motif **Ia** and motif **IV** preventing the protein from sampling **S1** when ATP is bound.

**Table 5.** Average separation distance and standard deviation between all residues in motif V with ATP and Mg^2+^ for states S1, S2, S3, and S4.

| Residues | Average Distance (Å) |  |  |  |
| --- | --- | --- | --- | --- |
|  | **S1** | **S2** | **S3** | **S4** |
| Val 533 | 5.3(8) | 4(1) | 3(1) | 4(2) |
| ASP 534 | 8(1) | 6(1) | 6.0(4) | 5(1) |
| SER 535 | 8(1) | 5(2) | 4.7(5) | 3(1) |
| SER 536 | 6.6(7) | 6.9(7) | 4.2(8) | 4.3(8) |
| GLN 537 | 9.2(8) | 8.5(8) | 7.2(5) | 6(1) |
| GLY 538 | 12(1) | 12.3(8) | 9(1) | 9(1) |
| SER 539 | 12.2(7) | 12.3(8) | 10(1) | 10.1(8) |
| GLU 540 | 11.0(4) | 11.0(7) | 10.0(6) | 10.1(5) |

These results are consistent with other studies in the literature that identify the importance of motif **V** in the communication between the ATP-pocket and the RNA-binding cleft of other SF1 and SF2 helicases. Molecular dynamics simulations of the flavivirus NS3 helicase protein revealed motif **V** as an allosteric link between the ATP-binding pocket and the RNA-binding cleft due to strong correlations between motif **V** and both binding pockets. ^51^ Furthermore, these results were supported by mutagenesis studies both *in vitro* and *in silico*.^52^ SARS-CoV-2 motif V and subdomain 2A are more dynamic than their flaviviral homologs but the mechanistic role of motif V is conserved. This further reflects the importance of motif V in the translocation mechanism making it an interesting target for antiviral development.

## Conclusions

To provide insight into the translocation mechanism utilized by nsp13 and the role ATP-binding plays in the translocation mechanism we performed simulations of Apo, ATP, ssRNA, and ssRNA+ATP ligand-bound states of the nsp13 helicase. Our models were verified by comparing substrate binding poses to crystal structure of other SF1 helicases that contain these substrates.

Interdomain distances revealed that the binding of ATP leads to an increase in the 1A–1B and 2A–1B domain distances corresponding to a widening of the RNA-binding cleft. A GMM-LDA approach revealed the presence of 4 states in the RNA-binding cleft. These four states represent a 4-step inchworm stepping translocation cycle where motif **Ia** and motif **IV** alternate in releasing a ssRNA phosphate before performing a powerstroke and binding a phosphate one basepair forward along ssRNA. The change in sampling of the four states in the ssRNA and ssRNA+ATP systems suggests that the first step in the cycle occurs due to the binding of ATP by nsp13. Analysis of the ATP-pocket of the 4 states reveal that motif **V** plays an important role in stabilizing ssRNA in states where motifs **Ia** and **IV** are largely separated. When ATP is present, motif **V** forms contacts with ATP reducing its interaction with ssRNA. Based on the simulations that we presented here motifs **Ia**, **IV**, and **V** play a crucial role in the translocation mechanism of nsp13. These motifs would be ideal targets for antiviral drugs to inhibit the function of nsp13.

Future work will focus on performing simulations of ssRNA+ADP+Pi and ss-RNA+ADP ligand-bound states of nsp13 to further investigate the translocation mechanism of nsp13 in the later stages of the hydrolysis cycle. Specifically, the role of ATP-hydrolysis, release of the inorganic phosphate, and release of ADP in the inchworm stepping mechanism will be elucidated. Also, enhanced sampling simulations of motif **V** would further clarify the ATP dependence of the structural conformations of motif **V** and its role in stabilizing the two phosphate gap in states where motif **Ia** and **IV** are largely separated.

## Supporting information

Parameter and starting files for simulation

## Supporting Information

Model verification data, ssRNA binding strength data, interdomain distance distributions, GMM score metrics, the specific residues used to calculated the distance between the motifs used in the LDA, LD eigenvectors for both the ATP-pocket and RNA-binding cleft LDA, and contacts between motif **V** and ssRNA phosphates.

## Acknowledgments

The computing for this project was performed at the High Performance Computing Center at Oklahoma State University supported in part through the National Science Foundation grant OAC-1531128. This work also used the Extreme Science and Engineering Discovery Environment (XSEDE), which is supported by National Science Foundation grant number ACI-1548562. We would specifically like to acknowledge the San Diego Supercomputer Center Comet used under XSEDE allocation CHE160008 awarded to MM. We would additionally like to thank Jake Anderson and Heidi Klem for their comments on the manuscript.

## References

1. WHO COVID-19 Explorer. Geneva: World Health Organization. 2020; https://worldhealthorg.shinyapps.io/covid/.

2. Adedeji, A. O.; Marchand, B.; te Velthuis, A. J. W.; Snijder, E. J.; Weiss, S.; Eoff, R. L.; Singh, K.; Sarafianos, S. G. Mechanism of nucleic acid unwinding by SARS-CoV helicase. PLoS One 2012, 7, e36521.

3. Kim, M. K.; Yu, M. S.; Park, H. R.; Kim, K. B.; Lee, C.; Cho, S. Y; Kang, J.; Yoon, H.; Kim, D. E.; Choo, H.; Jeong, Y. J.; Chong, Y. 2,6-Bis-arylmethyloxy-5-hydroxychromones with antiviral activity against both hepatitis C virus (HCV) and SARS-associated coronavirus (SCV). Eur. J. Med. Chem. 2011, 46, 5698–5704.

4. Adedeji, A. O.; Singh, K.; Calcaterra, N. E.; DeDiego, M. L.; Enjuanes, L.; Weiss, S.; Sarafianos, S. G. Severe acute respiratory syndrome coronavirus replication inhibitor that interferes with the nucleic acid unwinding of the viral helicase. Antimicrob. Agents Chemother. 2012, 56, 4718–4728.

5. Ndjomou, J.; Corby, M. J.; Sweeney, N. L.; Hanson, A. M.; Aydin, C.; Ali, A.; Schiffer, C. A.; Li, K.; Frankowski, K. J.; Schoenen, F. J.; Frick, D. N. Simultaneously Targeting the NS3 Protease and Helicase Activities for More Effective Hepatitis C Virus Therapy. ACS Chem. Biol. 2015, 10, 1887–1896.

6. Kumar, K.; Lupoli, T. J. Exploiting Existing Molecular Scaffolds for Long-Term COVID Treatment. ACS Med. Chem. Lett. 2020, 11, 1357–1360.

7. Romano, M.; Ruggiero, A.; Squeglia, F.; Maga, G.; Berisio, R. A Structural View of SARS-CoV-2 RNA Replication Machinery: RNA Synthesis, Proofreading and Final Capping. Cells 2020, 9.

8. Yan, L.; Zhang, Y.; Ge, J.; Zheng, L.; Gao, Y.; Wang, T.; Jia, Z.; Wang, H.; Huang, Y.; Li, M.; Wang, Q.; Rao, Z.; Lou, Z. Architecture of a SARS-CoV-2 mini replication and transcription complex. Nat. Commun. 2020, 11, 3–8.

9. Yan, L.; Ge, J.; Zheng, L.; Zhang, Y.; Gao, Y.; Wang, T.; Huang, Y.; Yang, Y.; Gao, S.; Li, M.; Liu, Z.; Wang, H.; Li, Y.; Chen, Y.; Guddat, L. W.; Wang, Q.; Rao, Z.; Lou, Z. Cryo-EM Structure of an Extended SARS-CoV-2 Replication and Transcription Complex Reveals an Intermediate State in Cap Synthesis. Cell 2021, 184, 184–193.e10.

10. Chen, J.; Malone, B.; Llewellyn, E.; Grasso, M.; Shelton, P. M.; Olinares, P. D. B.; Maruthi, K.; Eng, E. T.; Vatandaslar, H.; Chait, B. T.; Kapoor, T. M.; Darst, S. A.; Campbell, E. A. Structural Basis for Helicase-Polymerase Coupling in the SARS-CoV-2 Replication-Transcription Complex. Cell 2020, 182, 1560–1573.e13.

11. Ivanov, K. A.; Thiel, V.; Dobbe, J. C.; van der Meer, Y.; Snijder, E. J.; Ziebuhr, J. Multiple Enzymatic Activities Associated with Severe Acute Respiratory Syndrome Coronavirus Helicase. J. Virol. 2004, 78, 5619–5632.

12. Yuen, C. K.; Lam, J. Y.; Wong, W. M.; Mak, L. F.; Wang, X.; Chu, H.; Cai, J. P.; Jin, D. Y.; To, K. K. W.; Chan, J. F. W.; Yuen, K. Y.; Kok, K. H. SARS-CoV-2 nsp13, nsp14, nsp15 and orf6 function as potent interferon antagonists. Emerg. Microbes Infect. 2020, 9, 1418–1428.

13. Adedeji, A. O.; Singh, K.; Kassim, A.; Coleman, C. M.; Elliott, R.; Weiss, S. R.; Frieman, M. B.; Sarafianos, S. G. Evaluation of SSYA10-001 as a replication inhibitor of severe acute respiratory syndrome, mouse hepatitis, and Middle East respiratory syndrome coronaviruses. Antimicrob. Agents Chemother. 2014, 58, 4894–4898.

14. Kadaré, G.; Haenni, A. L. Virus-encoded RNA helicases. J. Virol. 1997, 71, 2583–2590.

15. Borowski, P.; Niebuhr, A.; Schmitz, H.; Hosmane, R. S.; Bretner, M.; Siwecka, M. A.; Kulikowski, T. NTPase/helicase of Flaviviridae: Inhibitors and inhibition of the enzyme. Acta Biochim. Pol. 2002, 49, 597–614.

16. Raney, K. D.; Sharma, S. D.; Moustafa, I. M.; Cameron, C. E. Hepatitis C virus non-structural protein 3 (HCV NS3): A multifunctional antiviral target. J. Biol. Chem. 2010, 285, 22725–22731.

17. Leung, D.; Schroder, K.; White, H.; Fang, N. X.; Stoermer, M. J.; Abbenante, G.; Martin, J. L.; Young, P. R.; Fairlie, D. P. Activity of Recombinant Dengue 2 Virus NS3 Protease in the Presence of a Truncated NS2B Co-factor, Small Peptide Substrates, and Inhibitors. J. Biol. Chem. 2001, 276, 45762–45771.

18. Byrd, C. M.; Grosenbach, D. W.; Berhanu, A.; Dai, D.; Jones, K. F.; Cardwell, K. B.; Schneider, C.; Yang, G.; Tyavanagimatt, S.; Harver, C.; Wineinger, K. A.; Page, J.; Stavale, E.; Stone, M. A.; Fuller, K. P.; Lovejoy, C.; Leeds, J. M.; Hruby, D. E.; Jordan, R. Novel benzoxazole inhibitor of dengue virus replication that targets the NS3 helicase. Antimicrob. Agents Chemother. 2013, 57, 1902–1912.

19. Sweeney, N. L.; Hanson, A. M.; Mukherjee, S.; Ndjomou, J.; Geiss, B. J.; Steel, J. J.; Frankowski, K. J.; Li, K.; Schoenen, F. J.; Frick, D. N. Benzothiazole and Pyrrolone Flavivirus Inhibitors Targeting the Viral Helicase. ACS Infect. Dis. 2015, 1, 140–148.

20. Lee, H.; Ren, J.; Nocadello, S.; Rice, A. J.; Ojeda, I.; Light, S.; Minasov, G.; Vargas, J.; Nagarathnam, D.; Anderson, W. F.; Johnson, M. E. Identification of novel small molecule inhibitors against NS2B/NS3 serine protease from Zika virus. Antiviral Res. 2017, 139, 49–58.

21. Mastrangelo, E.; Pezzullo, M.; De burghgraeve, T.; Kaptein, S.; Pastorino, B.; Dallmeier, K.; De lamballerie, X.; Neyts, J.; Hanson, A. M.; Frick, D. N.; Bolognesi, M.; Milani, M. Ivermectin is a potent inhibitor of flavivirus replication specifically targeting NS3 helicase activity: New prospects for an old drug. J. Antimicrob. Chemother. 2012, 67, 1884–1894.

22. Basavannacharya, C.; Vasudevan, S. G. Suramin inhibits helicase activity of NS3 protein of dengue virus in a fluorescence-based high throughput assay format. Biochem. Biophys. Res. Commun. 2014, 453, 539–544.

23. Shadrick, W. R.; Mukherjee, S.; Hanson, A. M.; Sweeney, N. L.; Frick, D. N. Aurintricarboxylic acid modulates the affinity of hepatitis C virus NS3 helicase for both nucleic acid and ATP. Biochemistry 2013, 52, 6151–6159.

24. Drummer, H. E.; Boo, I.; Maerz, A. L.; Poumbourios, P. A Conserved Gly436-Trp-Leu-Ala-Gly-Leu-Phe-Tyr Motif in Hepatitis C Virus Glycoprotein E2 Is a Determinant of CD81 Binding and Viral Entry. J. Virol. 2006, 80, 7844–7853.

25. Jia, Z.; Yan, L.; Ren, Z.; Wu, L.; Wang, J.; Guo, J.; Zheng, L.; Ming, Z.; Zhang, L.; Lou, Z.; Rao, Z. Delicate structural coordination of the Severe Acute Respiratory Syndrome coronavirus Nsp13 upon ATP hydrolysis. Nucleic Acids Res. 2019, 47, 6538–6550.

26. Jankowsky, E.; Fairman-Williams, M. E. RSC Biomol. Sci.; Royal Society of Chemistry, 2010; pp 1–31.

27. Fairman-Williams, M. E.; Guenther, U. P.; Jankowsky, E. SF1 and SF2 helicases: Family matters. Curr. Opin. Struct. Biol. 2010, 20, 313–324.

28. Ranji, A.; Boris-Lawrie, K. RNA helicases: Emerging roles in viral replication and the host innate response. 2010.

29. Pyle, A. M. RNA helicases and remodeling proteins. Curr. Opin. Chem. Biol. 2011, 15, 636–642.

30. Raney, K. D.; Byrd, A. K.; Aarattuthodiyil, S. Structure and mechanisms of SF1 DNA helicases. Adv. Exp. Med. Biol. 2013, 767, 17–46.

31. Seybert, A.; Hegyi, A.; Siddell, S. G.; Ziebuhr, J. The human coronavirus 229E superfamily 1 helicase has RNA and DNA duplex-unwinding activities with 5’-to-3’ polarity. Rna 2000, 6, 1056–1068.

32. Marchat, L. A.; Arzola-Rodríguez, S. I.; Hernandez-de la Cruz, O.; Lopez-Rosas, I.; Lopez-Camarillo, C. DEAD/DExH-Box RNA helicases in selected human parasites. Korean J. Parasitol. 2015, 53, 583–595.

33. Patel, S. S.; Donmez, I. Mechanisms of helicases. J. Biol. Chem. 2006, 281, 18265–18268.

34. Mickolajczyk, K. J.; Shelton, P. M.; Grasso, M.; Cao, X.; Warrington, S. E.; Aher, A.; Liu, S.; Kapoor, T. M. Force-dependent stimulation of RNA unwinding by SARS-CoV-2 nsp13 helicase. Biophys. J. 2021, 120, 1020–1030.

35. Miao, Y.; McCammon, J. A. Annu. Rep. Comput. Chem., 1st ed.; Elsevier B.V., 2017; Vol. 13; pp 231–278.

36. Chakrabarti, S.; Jayachandran, U.; Bonneau, F.; Fiorini, F.; Basquin, C.; Domcke, S.; Le Hir, H.; Conti, E. Molecular Mechanisms for the RNA-Dependent ATPase Activity of Upf1 and Its Regulation by Upf2. Mol. Cell 2011, 41, 693–703.

37. Theobald, D. L.; Wuttke, D. S. THESEUS: Maximum likelihood superpositioning and analysis of macromolecular structures. Bioinformatics 2006, 22, 2171–2172.

38. Cheng, Z.; Muhlrad, D.; Lim, M. K.; Parker, R.; Song, H. Structural and functional insights into the human Upf1 helicase core. EMBO J. 2007, 26, 253–264.

39. Case, D.; Betz, R.; Cerutti, D.; Cheatham III, T.; Darden, T.; Duke, R.; Giese, T.; Gohlke, H.; Goetz, A.; Homeyer, N.; Izadi, S.; Janowski, P.; Kaus, J.; Kovalenko, A.; Lee, T.; LeGrand, S.; Li, P.; Lin, C.; Luchko, T.; Luo, R.; Madej, B.; Mermelstein, D.; Merz, K.; Monard, G.; Nguyen, H.; Nguyen, H.; Omelyan, I.; Onufriev, A.; Roe, D.; Roitberg, A.; Sagui, C.; Simmerling, C.; Botello-Smith, W.; Swails, J.; Walker, R.; Wang, J.; Wolf, R.; Wu, X.; Xiao, L.; Kollman, P. AMBER 2018; University of California, San Francisco.: San Francisco, 2016.

40. Maier, J. A.; Martinez, C.; Kasavajhala, K.; Wickstrom, L.; Hauser, K. E.; Simmerling, C. ff14SB: Improving the Accuracy of Protein Side Chain and Backbone Parameters from ff99SB. J. Chem. Theory Comput. 2015, 11, 3696–3713.

41. Banáš, P.; Hollas, D.; Zgarbová, M.; Jurečka, P.; Orozco, M.; Cheatham, T. E.; Šponer, J.; Otyepka, M. Performance of molecular mechanics force fields for RNA simulations: Stability of UUCG and GNRA hairpins. J. Chem. Theory Comput. 2010, 6, 3836–3849.

42. Zgarbová, M.; Otyepka, M.; Šponer, J. J.; Mládek, A.; Banáš, P.; Cheatham, T. E.; Jurečka, P. Refinement of the Cornell et al. Nucleic Acids Force Field Based on Reference Quantum Chemical Calculations of Glycosidic Torsion Profiles. J. Chem. Theory Comput. 2011, 7, 2886–2902.

43. Meagher, K. L.; Redman, L. T.; Carlson, H. A. Development of polyphosphate parameters for use with the AMBER force field. J. Comput. Chem. 2003, 24, 1016–1025.

44. Li, P.; Merz, K. M. MCPB.py: A Python Based Metal Center Parameter Builder. J. Chem. Inf. Model. 2016, 56, 599–604.

45. Ryckaert, J. P.; Ciccotti, G.; Berendsen, H. J. Numerical integration of the cartesian equations of motion of a system with constraints: molecular dynamics of n-alkanes. J. Comput. Phys. 1977, 23, 327–341.

46. Darden, T.; York, D.; Pedersen, L. Particle mesh Ewald: An N·log(N) method for Ewald sums in large systems. J. Chem. Phys. 1993, 98, 10089–10092.

47. Pedregosa, F.; Varoquaux, G.; Gramfort, A.; Michel, V.; Thirion, B.; Grisel, O.; Blondel, M.; Prettenhofer, P.; Weiss, R.; Dubourg, V.; Vanderplas, J.; Passos, A.; Cournapeau, D.; Brucher, M.; Perrot, M.; Duchesnay, E. Scikit-learn: Machine Learning in Python. J. Mach. Learn. Res. 2011, 12, 2825–2830.

48. Hall, M. C.; Matson, S. W. Helicase motifs: The engine that powers DNA unwinding. Mol. Microbiol. 1999, 34, 867–877.

49. Wang, Q.; Arnold, J. J.; Uchida, A.; Raney, K. D.; Cameron, C. E. Phosphate release contributes to the rate-limiting step for unwinding by an RNA helicase. Nucleic Acids Res. 2009, 38, 1312–1324.

50. Garai, A.; Chowdhury, D.; Betterton, M. D. Two-state model for helicase translocation and unwinding of nucleic acids. Phys. Rev. E - Stat. Nonlinear, Soft Matter Phys. 2008, 77, 1–9.

51. Davidson, R. B.; Hendrix, J.; Geiss, B. J.; McCullagh, M. Allostery in the dengue virus NS3 helicase: Insights into the NTPase cycle from molecular simulations. PLoS Comput. Biol. 2018, 14, e1006103.

52. Du Pont, K. E.; Davidson, R. B.; McCullagh, M.; Geiss, B. J. Motif v regulates energy transduction between the flavivirus NS3 ATPase and RNA-binding cleft. J. Biol. Chem. 2020, 295, 1551–1564.

