## Supporting Information for "The Role of ATP in the RNA Translocation Mechanism of SARS-CoV-2 NSP13 Helicase"

### Supporting Information: The Role of ATP binding in the translocation mechanism of SARS-CoV-2 NSP13 Helicase.

### Model Verification

To verify our initial structures of the ATP, ssRNA, and ssRNA+ATP ligand-bound states of nsp13 we compare the contacts in our simulations to the contacts in the crystal structures of other SF1 helicases with ssRNA and ATP bound. The ssRNA contacts are shown in Table S1 and the ATP contacts are shown in Table S2.

**Table S1. Residues from motifs Ia, IV, and V in contact with RNA phosphates ($\leq$ 5.0 Å) for various SF1 RNA-bound helicase protein crystal structures and the percentage of frames where the corresponding nsp13 residues are in contact with RNA phosphates for both the RNA and RNA+ATP systems.**

| motif | Upf1 (2XZL)[1] | IGHMBP2 (4B3G)[2] | nsp13 | RNA | RNA+ATP |
|---|---|---|---|---|---|
| **Ia** | SER 461 | SER 244 | SER 310 | 83.15% | 84.25% |
| **Ia** | ASN 462 | ASN 245 | HIE 311 | 83.68% | 83.61% |
| **IV** | PRO 731 | PRO 540 | PRO 514 | 17.82% | 1.34% |
| **IV** | TYR 732 | TYR 541 | TYR 515 | 97.35% | 55.16% |
| **IV** | GLU 793 | ASN 542 | ASN 516 | 66.49% | 54.30% |
| **V** | SER 761 | SER 563 | SER 535 | 0.27% | 0.00% |
| **V** | ALA 764 | ASP 565 | VAL 533 | 44.14% | 0.12% |

**Table S2. Residues from motifs I, II, III, V and VI in contact with ATP or $MG^{2+}$ ($\leq$ 5.0 Å) for various SF1 ATP-bound helicase protein crystal structures and the percentage of frames where the corresponding nsp13 residues are in contact with ATP or $MG^{2+}$ for both the ATP and RNA+ATP systems.**

| motif | Upf1 (2GJK)[3] | nsp13 | ATP | RNA+ATP |
|---|---|---|---|---|
| **I** | GLY 492 | GLY 282 | 8.93% | 29.54% |
| **I** | PRO 493 | PRO 283 | 97.18% | 99.99% |
| **I** | PRO 494 | PRO 284 | 98.28% | 100.00% |
| **I** | GLY 495 | GLY 285 | 100.00% | 100.00% |
| **I** | THR 496 | THR 286 | 100.00% | 100.00% |
| **I** | GLY 497 | GLY 287 | 100.00% | 100.00% |
| **I** | LYS 498 | LYS 288 | 100.00% | 100.00% |
| **I** | THR 499 | SER 289 | 100.00% | 100.00% |
| **I** | VAL 500 | HIE 290 | 100.00% | 100.00% |
| **II** | ASP 636 | ASP 374 | 60.07% | 50.49% |
| **II** | GLU 637 | GLU 275 | 0.46% | 30.70% |
| **III** | GLN 665 | GLN 404 | 15.39% | 73.33% |
| **V** | GLY 831 | GLY 538 | 63.19% | 80.30% |
| **V** | ARG 832 | SER 539 | 36.77% | 19.22% |
| **V** | GLU 833 | GLU 540 | 99.27% | 84.46% |
| **VI** | ARG 865 | ARG 567 | 100.00% | 99.95% |
| **VI** | ARG 867 | LYS 569 | 80.17% | 45.12% |

### ssRNA Binding Strength

Inter-domain distance analysis of the Apo, ATP, ssRNA, and ssRNA+ATP states show that when nsp13 binds ATP there is a widening of the RNA-binding cleft. To measure the change in binding strength between nsp13 and ssRNA the linear interaction energy (Table S3) and root-mean-square fluctuation of the RNA phosphates (Table S4) are calculated for the ssRNA and ssRNA+ATP systems. The error in both analyses are too large to differentiate between the two systems. Figure S1 shows the labeling of the RNA phosphates and the highly conserved motifs of nsp13.

#### Linear Interaction Energy

**Table S3. Average linear interaction energy between each phosphate and protein residues within 12 Å for the ssRNA and ssRNA+ATP systems.**

| Linear Interaction Energy ($kcal \cdot mol^{-1}$) | | |
|---|---|---|
| Phosphates | ssRNA | ssRNA+ATP |
| P0 | -97(18) | -94(12) |
| P1 | -122(21) | -126(17) |
| P2 | -103(23) | -143(28) |
| P3 | -130(16) | -144(57) |
| P4 | -124(24) | -95(44) |
| P5 | -112(24) | -63(26) |
| P6 | -48(26) | -39(37) |

#### Fluctuations of ssRNA phosphates

**Table S4. RMSF of each phosphate for the ssRNA and ssRNA+ATP systems.**

| RMSF (Å) | | |
|---|---|---|
| Phosphates | ssRNA | ssRNA+ATP |
| P0 | 1.316(0.465) | 2.495(1.028) |
| P1 | 1.211(0.220) | 1.857(0.458) |
| P2 | 1.042(0.191) | 1.520(0.606) |
| P3 | 0.849(0.178) | 1.177(0.523) |
| P4 | 0.879(0.205) | 1.012(0.038) |
| P5 | 1.328(0.268) | 1.322(0.214) |
| P6 | 2.427(0.812) | 2.204(0.994) |

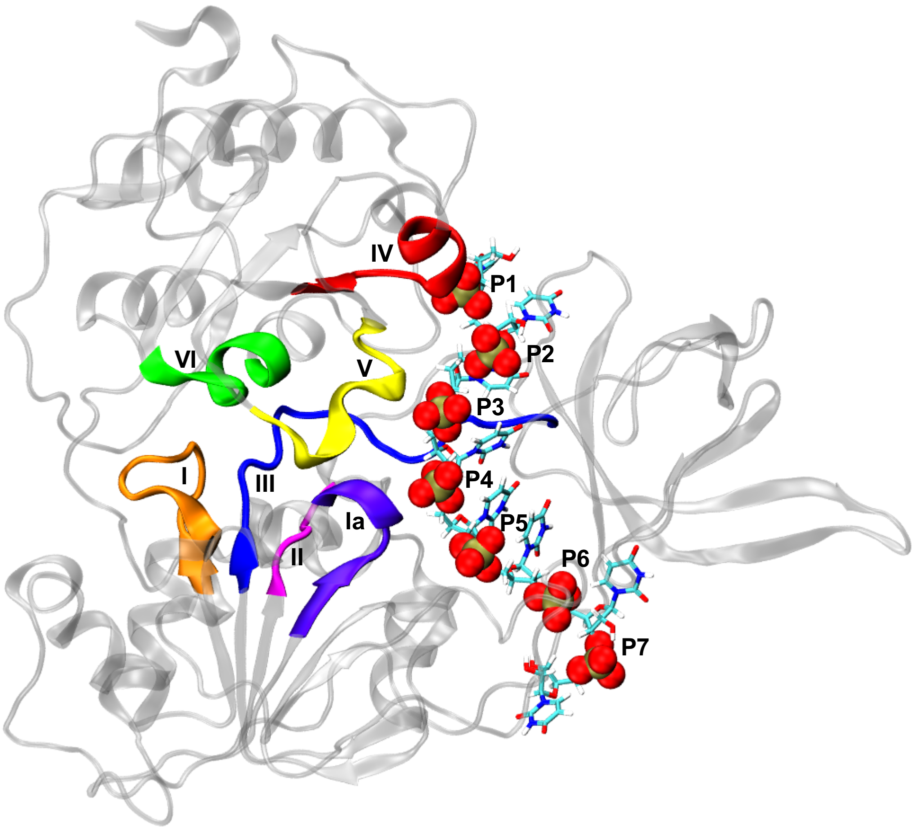


**Figure S1.** Representative structure of nsp13 with ssRNA bound. Motifs **I** (orange), **Ia** (violet), **II** (magenta), **III** (blue), **IV** (red), **V** (yellow), **VI** (green), and each phosphate in the ssRNA backbone is highlighted and labeled. The Zinc-Binding domain and Stalk domain are removed for clarity.

### Inter-domain Distances

The inter-domain distances between domains 1A, 2A, and 1B were calculated for the Apo, ATP, ssRNA, and ssRNA+ATP ligand-bound states of nsp13. The distributions of the 1A–1B, 2A–1B, and 1A–2A distances for each ligand-bound state are shown in Figure S2(a-c), respectively.

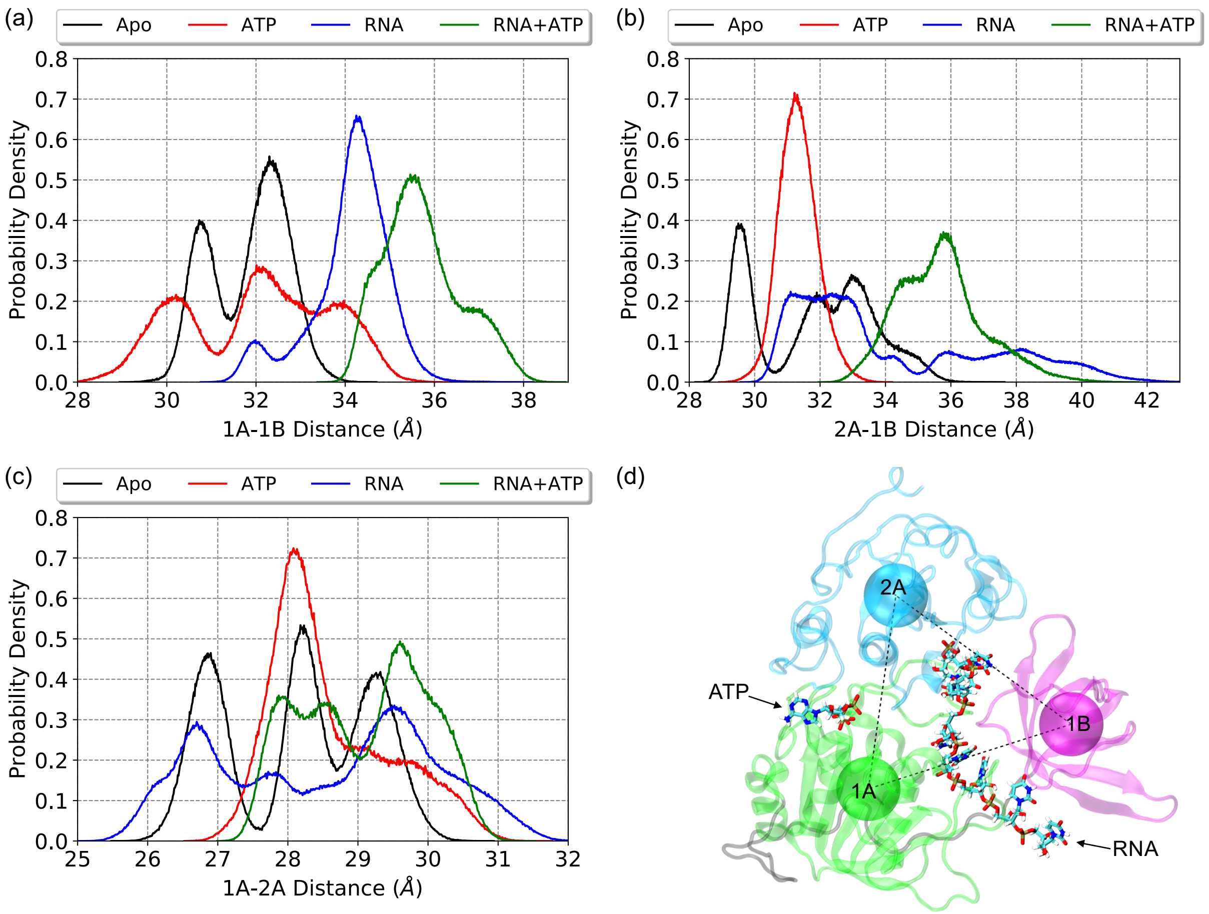


**Figure S2.** Probability density of the center-of-mass separation distance between domains (a) 1A–1B, (b) 1A–2A, and (c) 2A–1B of the nsp13 Apo, ATP, ssRNA, and ssRNA+ATP ligand-bound states. (d) Structural depiction of the center-of-mass of domains 1B (magenta), 1A (green), and 2A (cyan)

### Gaussian Mixture Model and Linear Discriminant Analysis

Figure S3 shows the Silhouette, Calinski-Harabasz (CH), and Davies-Bouldin (DB) scores for cluster sizes ranging from two clusters to ten clusters for the RNA-binding cleft distances. Based on the maximums of the Silhouette and CH scores and minimums of the DB score a cluster size of four was chosen. Linear discriminant analysis (LDA) was utilized to differentiate between the 4 states in the RNA-binding cleft and the ATP-pocket. Table S5 shows the $\alpha$-carbon of the residues used to represent the position of each motif used in the LDA. The LD1 and LD2 vectors for the RNA-binding cleft and the ATP pocket are shown in Table S6 and Table S7, respectively.

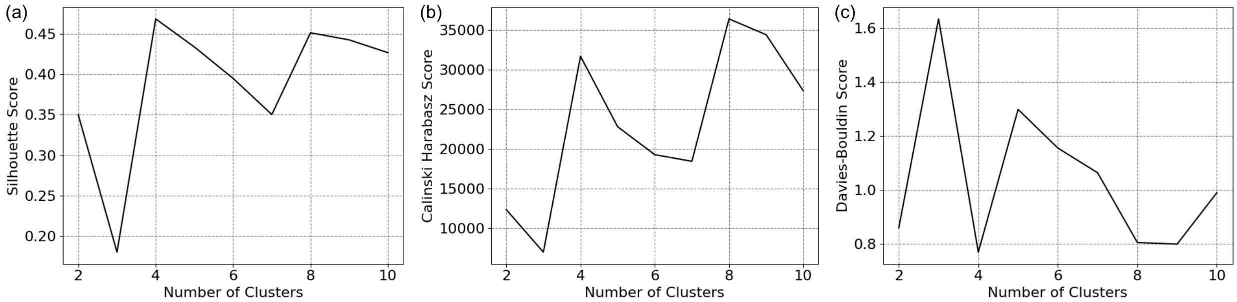


**Figure S3.** (a) Silhouette, (b) Calinski-Harabasz, and (c) Davies-Bouldin scores for various cluster sizes from GMM clustering of the RNA-binding cleft.

**Table S5. Residues used as the position of each motif utilized by the linear discriminant analysis in calculating the difference between states S1, S2, S3, and S4.**

| Motif | Residue |
|---|---|
| **I** | GLN 281 |
| **Ia** | HID 311 |
| **II** | ILE 375 |
| **IV** | ASN 516 |
| **V** | ASP 534 |
| **VI** | ARG 567 |

**Table S6. Coefficients for each distance used in the linear discriminant analysis to describe the RNA-binding cleft for LD1 and LD2.**

| LDA Coefficients | | |
|---|---|---|
| Residues | LD1 | LD2 |
| **IV – P** | -0.424 | 0.335 |
| **IV – Ia** | -0.095 | -0.657 |
| **Ia – P** | 1.859 | -0.050 |

**Table S7. Coefficients for each distance used in the linear discriminant analysis to describe the ATP-pocket for LD1 and LD2.**

| LDA Coefficients | | |
|---|---|---|
| Residues | LD1 | LD2 |
| **I – V** | 0.931 | -1.080 |
| **I – Ia** | -1.173 | 1.427 |
| **Ia – V** | -0.520 | -0.318 |
| **II – V** | -0.765 | 0.207 |
| **II – VI** | -0.096 | 0.020 |
| **IV – V** | -0.373 | -0.067 |
| **V – VI** | -1.024 | 0.509 |
| **V – P** | 0.646 | 0.244 |

### Motif V–ssRNA Phosphate Contacts

Table S8 shows the percentage of frames that motif **V** was in contact with each ssRNA phosphate. If any residue of motif **V** was within 5 Å of an ssRNA phosphate than it was considered a contact. The phosphates are labeled relative to the phosphate bound by motif **Ia**. Table S9 shows the average separation distance between each residue in motif **V** and the closest ssRNA phosphate.

**Table S8. Percentage of frames where motif V is bound ($\leq$ 5.0 Å) to each ssRNA phosphates. Phosphates are labeled relative to the phosphate motif Ia is binding, where motif Ia is binding $P_n$.**

| Residues | **S1** | **S2** | **S3** | **S4** |
|---|---|---|---|---|
| $P_n$ | 0.88% | 7.02% | 0.10% | 0.46% |
| $P_{n-1}$ | 53.60% | 44.14% | 62.50% | 77.07% |
| $P_{n-2}$ | 43.92% | 0.39% | 1.87% | 12.35% |
| $P_{n-3}$ | 0.56% | 0.00% | 0.00% | 0.01% |

**Table S9. Average separation distance and standard deviation between all residues in motif V with ssRNA phosphates for states S1, S2, S3, and S4.**

| | Average Distance (Å) | | | |
|---|---|---|---|---|
| Residues | **S1** | **S2** | **S3** | **S4** |
| Val 533 | 16(2) | 19.2(8) | 18.6(6) | 17(2) |
| ASP 534 | 12(2) | 15(1) | 14.6(7) | 13(2) |
| SER 535 | 11(1) | 15(1) | 12.7(8) | 12(2) |
| SER 536 | 6(2) | 10(1) | 9.9(9) | 8(1) |
| GLN 537 | 7(1) | 10.8(6) | 10.1(5) | 8(2) |
| GLY 538 | 4(2) | 7.9(8) | 7.4(7) | 6(3) |
| SER 539 | 4.3(6) | 5.1(6) | 4.8(5) | 4.6(9) |
| GLU 540 | 7.1(9) | 8.6(5) | 7.9(9) | 7(2) |

### References


1. Chakrabarti, S.; Jayachandran, U.; Bonneau, F.; Fiorini, F.; Basquin, C.; Domcke, S.; Le Hir, H.; Conti, E. Molecular Mechanisms for the RNA-Dependent ATPase Activity of Upf1 and Its Regulation by Upf2. *Mol. Cell* **2011**, *41*, 693–703.
2. Lim, S. C.; Bowler, M. W.; Lai, T. F.; Song, H. The Ighmbp2 helicase structure reveals the molecular basis for disease-causing mutations in DMSA1. *Nucleic Acids Res.* **2012**, *40*, 11009–11022.
3. Cheng, Z.; Muhlrad, D.; Lim, M. K.; Parker, R.; Song, H. Structural and functional insights into the human Upf1 helicase core. *EMBO J.* **2007**, *26*, 253–264.
